# PeruNPDB: The Peruvian Natural Products Database for in silico drug screening

**DOI:** 10.1101/2023.01.15.524152

**Authors:** Haruna Luz Barazorda-Ccahuana, Lena Gálvez Ranilla, Mayron Antonio Candia-Puma, Eymi Gladys Cárcamo-Rodriguez, Angela Emperatriz Centeno-Lopez, Gonzalo Davila Del-Carpio, José L. Medina-Franco, Miguel Angel Chávez-Fumagalli

## Abstract

Since the number of drugs based on natural products (NPs) represents a large source of novel pharmacological entities, NPs have acquired significance in drug discovery. Peru is considered a megadiverse country with many endemic species of plants, terrestrial, and marine animals, and microorganisms. NPs databases have a major impact on drug discovery development. For this reason, several countries such as Mexico, Brazil, India, and China have initiatives to assemble and maintain NPs databases that are representative of their diversity and ethnopharmacological usage. We describe the assembly, curation, and full chemoinformatic evaluation of the content and coverage in chemical space, as well as the physicochemical attributes and chemical diversity of the initial version of the Peruvian Natural Products Database (PeruNPDB), which contains 280 compounds. Access to PeruNPDB is available for free (https://perunpdb.com.pe/). The PeruNPDB’s collection is intended to be used in a variety of tasks, such as virtual screening campaigns against various disease targets or biological endpoints. This emphasizes the significance of biodiversity protection both directly and indirectly on human health.

## Introduction

Biodiversity is the variety of all life forms, including the morphological diversity of individuals and populations within a species, the taxonomic diversity of species within a community or ecosystem, the functional diversity of groups of species within an ecosystem, and the diversity of ecosystems themselves. [1]. While the total number of species in every taxonomic group has been predicted for all kingdoms of life on earth at approximately 8.7 million [2,3], it is remarkable that the distribution of that vast number of species is highly concentrated in specific areas. These regions are particularly important for biodiversity conservation and are called biodiversity hotspots [4], although: Bolivia, Brazil, China, Colombia, Costa Rica, the Democratic Republic of Congo, Ecuador, India, Indonesia, Kenya, Madagascar, Malaysia, Mexico, Peru, Philippines, South Africa, and Venezuela are considered megadiverse countries [5]. Peru occupies the seventh place in this group, as it possesses 28 of the 32 existing climates in the world and 84 of the 103 life zones known on earth. This is evidenced by considering that the country has 25,000 plant species or 10% of the entire number of species worldwide, whereas 30% are endemic, and endemic animal species such as 115 birds, 109 mammals, and 185 amphibians species, which represent 6, 27.5 and 48.5% of the total number worldwide, respectively [6,7].

Biodiversity conservation is important since plants, animals, and other life forms such as bacteria, archaea, protozoa, and fungi, are used directly or indirectly to produce pharmaceuticals, and for their scientific value, among other resources [8]. The number of drugs derived from natural products (NP) that were introduced to the market over forty years represented a significant source of new pharmacological entities. [9]. Whilst the Peruvian population uses approximately 5000 Peruvian plants for 49 purposes or applications, where about 1400 species are described as medicinal [10–13]. The contribution from traditional Peruvian medicine can be embodied by Quinine, a component of the bark of the cinchona tree (*Cinchona officinalis*), employed in the treatment of malaria [14]. Additionally, two other valuable contributions to modern pharmacopeias such as the coca plant (*Erythroxylum coca*), from which cocaine was first isolated and later led to local anesthetics [15], and the balsam of Peru (*Myroxylon balsamum*), which was used wide-reaching for the treatment of wounds [16], can be mentioned. However, the potential of Peruvian NPs remains underexploited since most of these useful native species can be domesticated or semi-domesticated [17]. Also, the amount and nature of experimental evidence published on active compounds are still limited [18], and most of the current studies reported crude medicinal activities, while potentially active compounds have been isolated only from a few numbers of plants [19].

Computer-aided drug design (CADD), one of the key approaches to modern pre-clinical drug discovery, can be defined as computational methods that are applied to discover, develop, and analyze drugs and active molecules [20]. Among the key approaches that comprise CADD, virtual screening is one of the major contributors to CADD since it stands as a contemporary approach to the experimental in vitro high-throughput screening (HTS) for hit identification and optimization [21]. Integrating CADD approaches to curated databases, which are described as a well-organized collection of data in any field, the drug development process may be sped up and cost reduced.[22]. Considering this, large databases containing NPs from various data sources have been released, such as the COlleCtion of Open Natural prodUcTs (COCONUT), which contains 406,076 unique “flat” NPs, and a total of 730,441 NPs where stereochemistry has been preserved [23]; and the LOTUS initiative, which has 750,000 referenced structure-organism pairs [24]. Also, several NPs compound databases from particular geographical locations have been assembled, such as the Traditional Chinese Medicine (TCM) Database@Taiwan database containing approximately 58,000 molecules [25]; the Indian Medicinal Plants, Phytochemistry and Therapeutics 2.0 (IMPPAT 2.0) which contains more than 10,000 phytochemicals [26]; and the AfroDB which is composed of around 1000 NPs [27]. Likewise, some countries in Latin America have published their own public NPs databases such as the NuBBE_DB_ from Brazil which contains more than 2000 NPs [28], and BIOFACQUIM from Mexico, which contains a total of 531 molecules [29]. Furthermore, NPs databases had been used as a repository to identify several promising candidates to be considered for further development for the treatment of diseases [30], such as Chagas disease [31,32], Tuberculosis [33], Leishmaniasis [34,35], Schistosomiasis [36], and COVID-19 [37]. The present work introduces the first version of the Peruvian Natural Products Database (PeruNPDB), describing its assembly, curation, and chemoinformatic characterization of molecular diversity and coverage in chemical space. The database is freely available at the web-interface PeruNPDB Explorer (https://perunpdb.com.pe/). We anticipate that the PeruNPDB will make it possible to conduct additional virtual screening tests to create innovative pharmacological entities and other biotechnological approaches and serve as a resource for information on conservation guidelines.

## Methods

### Search strategy, study selection, and data extraction

A systematic review search strategy to examine the literature for studies describing compounds from Peruvian NPs was adapted from [38]. Whereas PubMed, the main database for the health sciences, maintained by the National Center for Biotechnology Information (NCBI) at the National Library of Medicine (NLM), is a database that contains about 32 million citations, belonging to more than 5,300 journals currently indexed in MEDLINE [39]; it provides uniform indexing of biomedical literature, the Medical Subject Headings (MeSH terms), which form a controlled vocabulary or specific set of terms that describe the topic of a paper consistently and uniformly [40]. Firstly, to find terms associated in the literature with Peruvian NPs, the MeSH terms “*Peru*” AND the “*Natural Products*” were employed in a search carried out at the PubMed database (https://pubmed.ncbi.nlm.nih.gov/), (last searched on 10 June 2022), though the results were plotted into a network map of the co-occurrence of MeSH terms in the VOSviewer software (version 1.6.17) [41], which employs a modularity-based method algorithm to measure the strength of clusters [42]. The resultant cluster content was analyzed to select relevant studies associated with Peruvian NPs.

Three phases went into selecting the studies. First, papers written in languages other than English, copies of articles, reviews, and meta-analyses were disregarded. The highly relevant full studies were then retrieved and separated from the papers with a title or abstract that did not provide enough information to be included. Next, the titles and abstracts of the publications chosen through the search approach were visually evaluated. The data supplied from each investigation contained the compounds’ characterization as well as details on the genus and species of the NPs from which the compounds were isolated. Additionally, the information from the bibliographic reference was extracted, even if all research that discussed chemicals derived from Peruvian natural sources was already considered.

### PeruNPDB assemble and molecular properties calculation

The simplified molecular-input line-entry system (SMILES) [43] of compounds previously described in the NPs selected in the previous step were searched and retrieved from PubChem [44], DrugBank [45], or ChEMBL [46] servers, while for unavailable compounds the ChemDraw tool [47] was employed to generate the SMILE notation. Moreover, the Osiris DataWarrior v05.02.01 software [48] was employed to generate the dataset’s structure data files (SDFs). This followed the uploading to the Konstanz information miner (KNIME) Analytics Platform [49], where the “Molecular Type Cast”, and the “RDKit Structure Normalizer” KNIME nodes were employed to curate the chemical structures on the dataset. Moreover, for every compound in the dataset, the classification system for describing small molecule structures is described based on NP Classifier [50], which employs a biosynthetic ontology that is specific to natural products; or ClassyFire [51] which is a general classification system for small molecules that are based on the ChemOnt ontology, was employed.

The KNIME’s “RKDit Descriptor Calculator” node was employed to calculate six physicochemical properties of therapeutic interest, namely: molecular weight (MW), octanol/water partition coefficient (clogP), topological surface area (TPSA), aqueous solubility (clogS), number of H-bond donor atoms (HBD) and number of H-bond acceptor atoms (HBA) of the PeruNPDB, while the statistical analysis was done within the GraphPad Prism software version 9.4.0 (673) for Windows, GraphPad Software, San Diego, California USA, www.graphpad.com, by calculating the mean, median, standard deviation, and the coefficient of variation of the calculated properties. Box-and-whisker plots showing, the maximum and minimum values were generated for visualization, and the One-way ANOVA followed by Dunnett correction for multiple comparisons test was employed to evaluate the differences between the datasets. The results were considered statistically significant when p<0.05.

### Visual representation of chemical space

To generate a visual representation of the chemical space of the PeruNPDB, two visualization methods were employed: principal component analysis (PCA), which reduces data dimensions by geometrically projecting them onto lower dimensions called principal components (PCs) [52] calculated by the “PCA” KNIME node. The second technique was the t-distributed stochastic neighbor embedding (t-SNE), which is a nonlinear dimension reduction in which Gaussian probability distributions over high-dimensional space are constructed and used to optimize a student t-distribution in low-dimensional space [53], calculated by the t-SNE (L. Johnson)” KNIME node. Three and two-dimensional scatter-plot representations were generated for PCA and t-SNE, respectively with the Plotly KINME node [54]. Additionally, the Tanimoto similarity score was calculated for clustering the compounds, while the atom-pair-based fingerprints of the compounds were obtained using the “ChemmineR” package [55] in the R programming environment (version 4.0.3) [56], a heatmap was generated for visualization. The same procedure was employed in the reference datasets: AfroDB [27], BIOFAQUIM [29], and NUBBE_DB_ [28] retrieved from the ZINC20 database [57].

### Global diversity: consensus diversity analysis

Since chemical diversity strongly depends on the structure representation, it is reasonable to consider multiple representations for a complete global assessment. The consensus diversity (CD) plots have been proposed as simple two-dimensional graphs that enable the comparison of the diversity of compound data sets using four sets of structural representations: the molecular fingerprints, scaffolds, molecular properties, and the number of compounds [58]. The multiple-variable plot was generated by GraphPad Prism software version 9.4.0 (673), whereas the y-axis represents the area under the cyclic system recovery curve [59], the x-axis, represents the median of the fingerprint-based diversity computed with Molecular Access System (MACCS) keys (166-bits) and the Tanimoto coefficient [60], the bubble color represents the molecular properties of pharmaceutical interest, and the bubble size represents the number of compounds for each database.

### Drug-likeness

The Osiris DataWarrior v05.02.01 software [61] was employed to calculate the drug-likeness score of the compounds from the PeruNPDB; the calculation is based on a library of ~5300 substructure fragments and their associated drug-likeness scores. This library was prepared by fragmenting 3,300 commercial drugs as well as 15,000 commercial non-drug-like Fluka compounds [61]. Frequency distribution of the obtained scores was performed at GraphPad Prism software version 9.4.0 (673) for Windows, GraphPad Software, San Diego, California USA (www.graphpad.com), and plotted into stacked bar plots. Furthermore, the Lipinski Rule-of-5 (Ro5) is a set of four rules (logP, MW, and H-bond donor and acceptor cut-offs) for drug-likeness and oral bioavailability derived from a subset of 2,245 drugs [62]. For this Lipinski’s Ro5 KNIME node was employed to assess the number of violations to the rule for each compound on the PeruNPDB and plotted into pie charts. The US Food and Drug Administration (FDA)-approved drugs dataset [57], was employed as a reference, whereas the same procedures were applied to their compounds. Also, the chemical space representation was analyzed, and the procedures were the same as described earlier.

## Results

### PeruNPDB assemble

In the present study, the assembly of the PeruNPDB, followed by its chemoinformatic characterization on molecular diversity and coverage of the chemical space was performed; to select the studies from which the compounds will further retrieve, a search with the MeSH Terms “*Peru*” AND “*Natural Products*” was performed in the Pubmed database, followed by the construction of a network map of the co-occurrence of MeSH terms. The search resulted in 399 published papers between 1950–2021-, whereas establishing the value of five as the minimum number of occurrences of keywords, a map with 194 keywords that reaches the threshold was constructed (Figure 1A). In the analysis of the map, it is shown that six main clusters were formed, while terms such as “*Plant Extracts*”, “*Plants, medicinal*”, “*Phytotherapy*", “*Ethnopharmacology*", “*Ethnobotany*", “*Plants stems*", “*Plants bark*", and “*Seeds*”, which are associated with NPs were observed in the first cluster (red color). Also, terms such as “*Peru*”, “*Humans*”, “*Animals*”, and “*Male*”, were recurrent terms. Although using the eligibility criterion established, 47 articles were selected which showed a 2000–2021-year range, and terms such as “*Flavonoids*”, “*Sesquiterpenes*”, and “*Anthocyanins*”, were recurrent terms (Figure 1B). Also, bibliographic data extracted from the selected articles analyzed: the “*Journal of Agricultural and Food Chemistry*”, the “*Journal of Ethnopharmacology*”, “*Phytochemistry*”, and “*Planta Medica*” where the main peer-reviewed journals were the studies describing compounds extracted from Peruvian NPs were published (Figure 1C).

**Figure 1.**
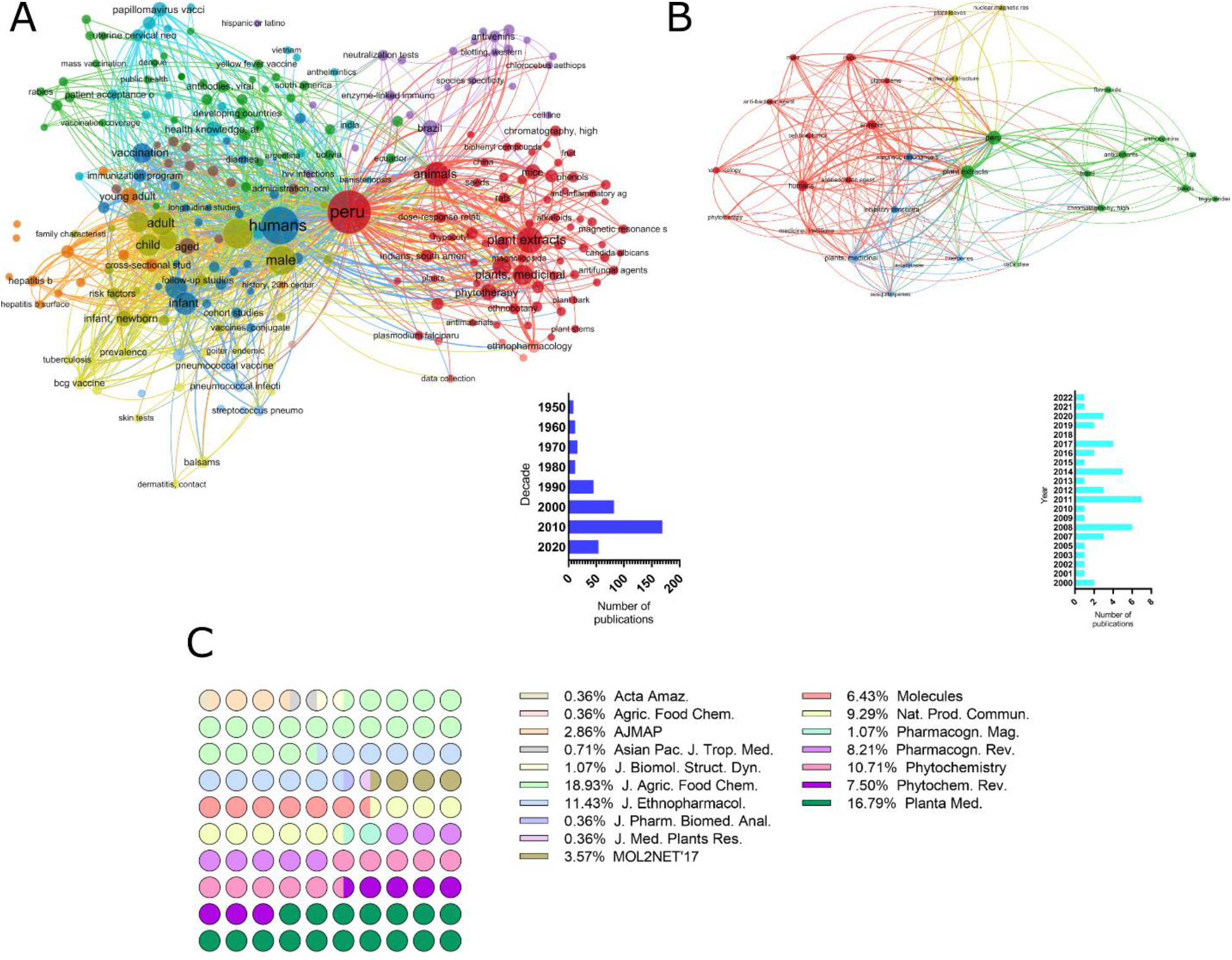
Bibliographic search for studies describing the characterization of Peruvian natural products. A. Network map of the co-occurrence of MeSH terms. B. Network map of articles selected from 2000–2021-year. C. Bibliographic data extracted from selected articles that describe compounds extracted from Peruvian NPs.

Furthermore, while retrieving the SMILES of the compounds from PubChem, DrugBank, and ChEMBL, it was observed that 242 structures were found in the repositories and that 38 needed to be generated in the ChemDraw tool. Ninety-five and five percent of the compounds were retrieved from plant or animal sources, respectively (Figure 2A). The genus from which most of the compounds were extracted were Unicaria and Lepidium, with 11 and 10 percent, respectively (Figure 2B). When analyzing the structure of the compounds with a classification system for small molecule structures, it is shown that 76 classes of NPs were found among the 280 compounds of the PeruNPDB, whereas anthocyanidins (N=25), aporphine alkaloids (N=11), cinnamic acids and derivatives (N=17), germacrane sesquiterpenoids (N=13), stigmastane steroids (N=10), and unsaturated fatty acids(N=22) were the most predicted classes of compounds (Figure 3).

**Figure 2.**
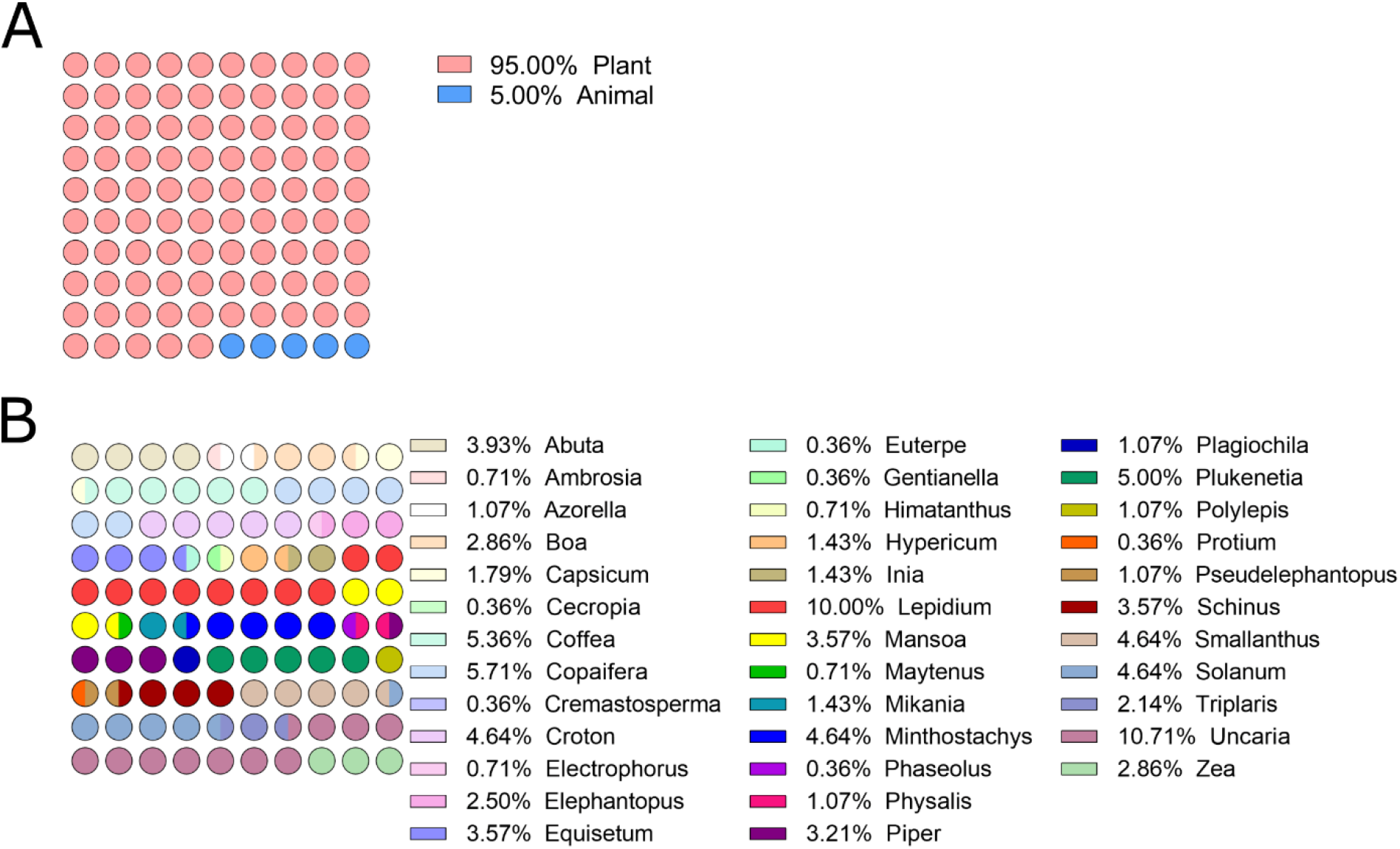
Dot plots showing the kingdom and genus of the species studied. A. Compounds of Peruvian NPs found in PubChem, DrugBank, and ChEMBL databases. B. Dot plot of the genus of the Peruvian NPs compounds obtained from the databases.

**Figure 3.**
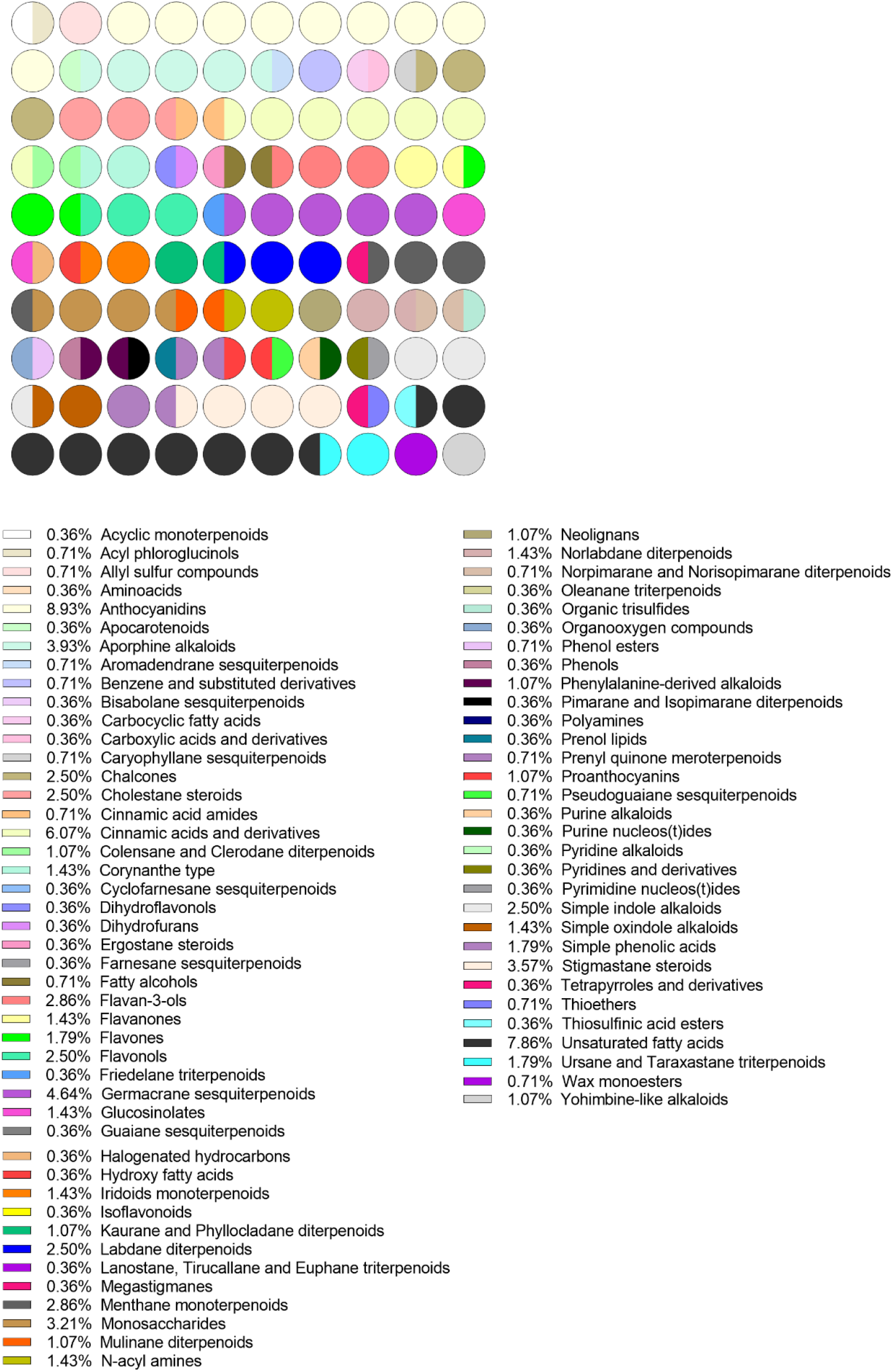
Dot plots showing the natural products classification.

### Molecular properties

Six physicochemical properties were calculated for all compounds in PeruNPDB and plotted into box plots, which include the distribution of the same properties of the three reference datasets, retrieved from the ZINC20 database (Figure 4). To compare the results of the datasets, the coefficient of variation (CV) was calculated. which represents the ratio of the standard deviation to the mean and is considered a useful tool to statistically compare the degree of variation from one dataset to another [53]. Besides the results of the HBA, in which NuBBE_DB_ obtain the highest CV (123.2%), the PeruNPDB showed the highest CV in MW, clogP, TPSA, clogS, and HBD with 46.58, 84.49, 112.8, 50.08 and 83.84%, respectively. Still, the results from TPSA, clogP, clogS, and HBD showed high statistical differences compared to AfroDB, BIOFAQUIM, and NuBBE_DB_, while showed no statistical difference in HBA results compared to the AfroNP database (Figure 4).

**Figure 4.**
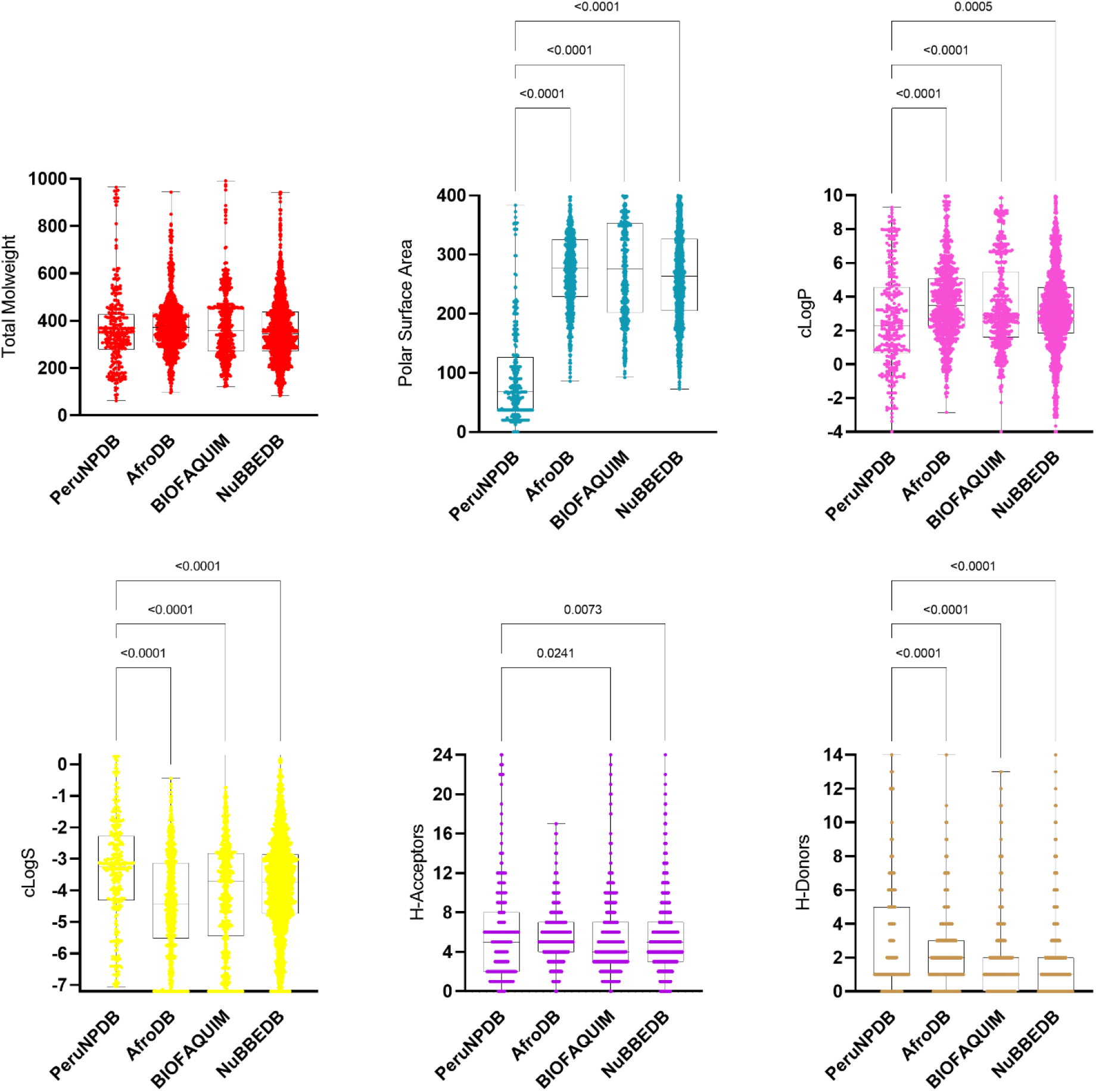
Box plots for the physicochemical properties of PeruNPDB and reference datasets.

### Visualization of the chemical space

The chemical space visualization of PeruNPDB was conducted using PCA t-SNE. Though the visual analysis of 3D-PCA shows that molecules in PeruNPDB share the chemical space roughly with NuBBE_DB_ Whereas in some regions the molecules of PeruNPDB are predominant (Figure 5A). PeruNPDB, BIOFAQUIM, and NuBBE_DB_ chemicals overlap in most of the chemical space represented, according to the 2D-t-SNE visual analysis (Figure 5B).

**Figure 5.**
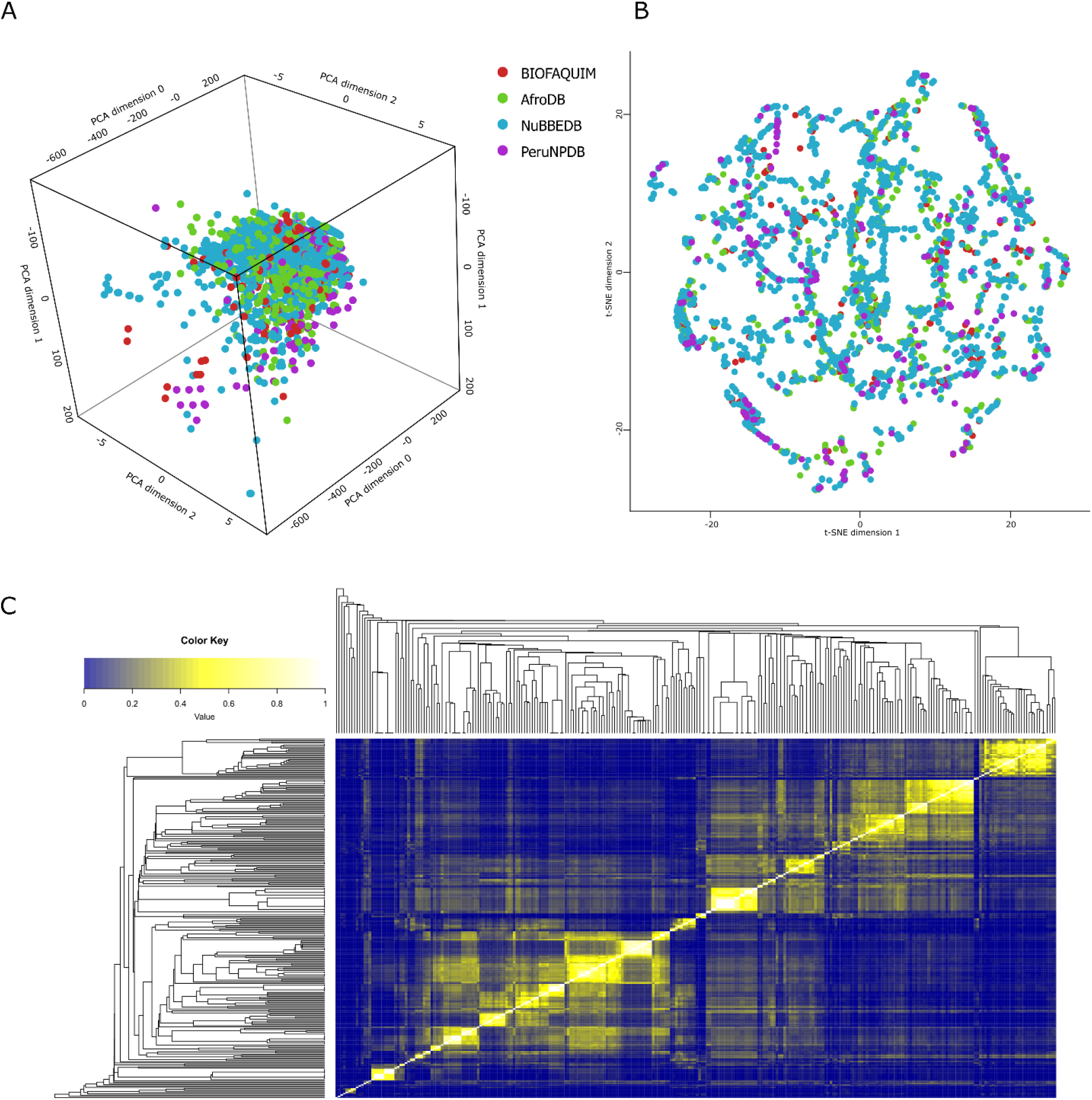
Visual representation of the chemical space of the PeruNPDB and reference datasets. A. PeruNPDB 3D-PCA chemical space. B. 2D-t-SNE visual analysis of the compounds PeruNPDB, AfroNP, BIOFAQUIM, and NuBBEDB. C. Heatmap generated with Tanimoto scoring matrix of similar structures among compounds between PeruNPDB and control data sets.

### Diversity analysis

The heatmap generated using the Tanimoto score matrix and the atom-pair-based fingerprints show that there is a similarity between the structures of the compounds of the PeruNPDB, AfroDB, BIOFAQUIM, and NuBBE_DB_ (Figure 5C). Also, a consensus diversity plot was used to evaluate the diversity of the PeruNPDB dataset, based on molecular fingerprints, scaffolds, and physicochemical properties. The Euclidean distance of the scaled properties was used to compute the property-based diversity of the PeruNPDB, AfroDB, BIOFAQUIM, and NuBBE_DB_ databases. Data points on a continuous color scale are used to represent the values on the color CD plot. Darker colors signify less diversity, but brighter colors signify more diversity. Finally, different point sizes are used to illustrate how large or tiny the databases are, with smaller data points indicating databases with fewer molecules. The results showed that the diversity of compounds found in the PeruNPDB was the largest since it was found in the area where the highest diversity in scaffold and fingerprints should are found (Figure 6), which is consistent with the results shown in the box plots (Figure 5).

**Figure 6.**
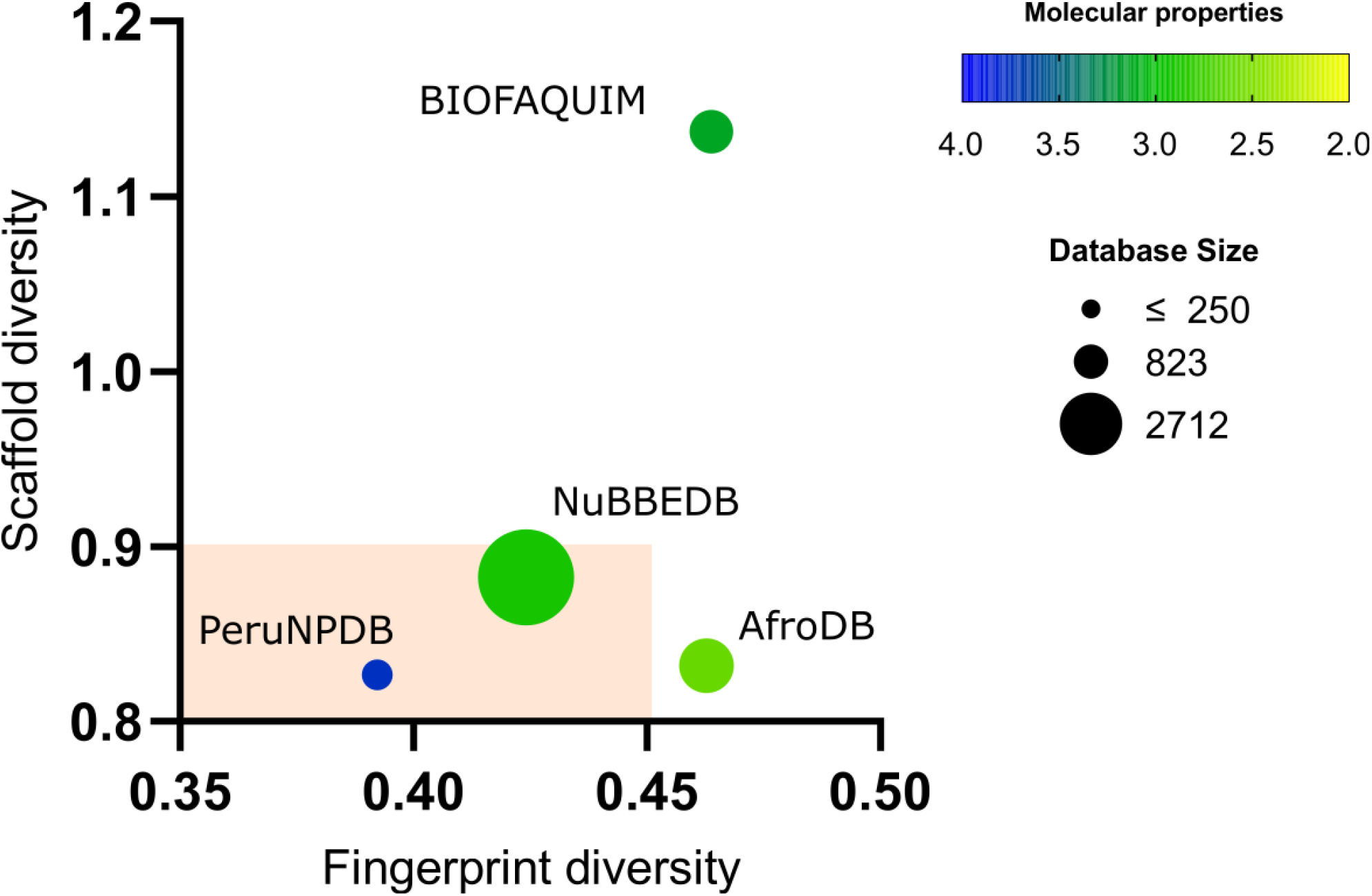
Consensus Diversity Plot comparing the global diversity of PeruNPDB with the reference data sets.

### Drug-likeness

Druglikeness assesses qualitatively the chance for a molecule to become an oral drug concerning bioavailability and is established from structural or physicochemical inspections of development compounds advanced enough to be considered oral drug candidates [63]. To assess the “drug-like” profile of the compounds from the PeruNPDB two approaches were performed; firstly, the frequency distribution of the drug-likeness score was analyzed, and the results showed that besides the differences in the number of compounds compared in both datasets a similar distribution is observed among the compounds is observed (Figure 7A)., In the second approach, the number of violations to Lipinski’s Ro5 was analyzed and the results showed that compounds with at least one violation represent the 85.82 and 76.35% of the FDA and PeruNPDB datasets, respectively (Figure 7B). Also, the visual representation of the chemical space as PCAs (Figure 7C) and t-SNE (Figure 7D) indicates that some of the NPs are distributed in the same space as the already approved drugs. The findings imply that because the compounds in PeruNPDB have chemical structures like those of approved medications, they can be used in virtual screening to find possible lead compounds or points for further optimization.

**Figure 7.**
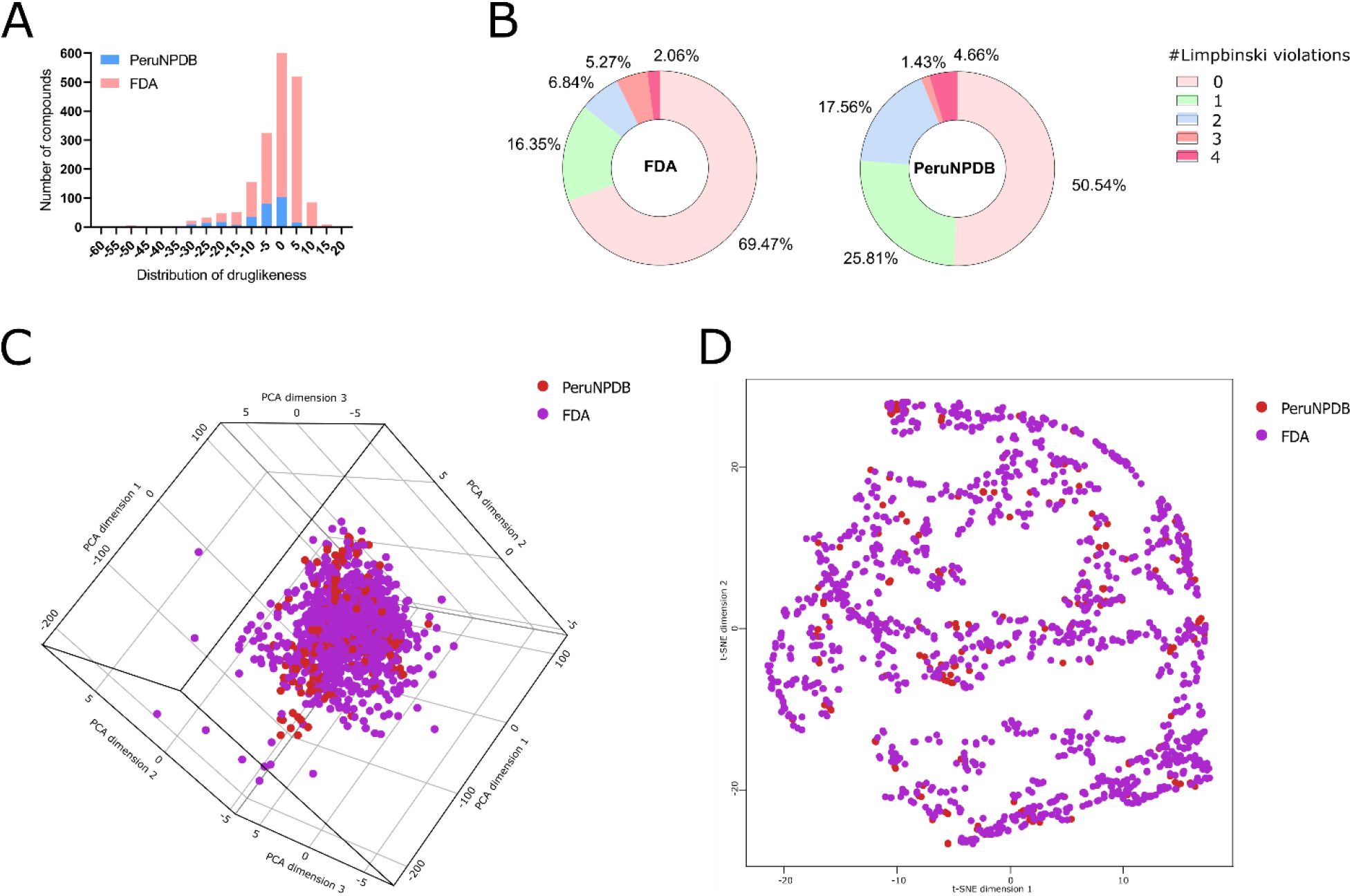
Druglikeness analysis of the PeruNPDB and the reference datasets. A. The similar distribution between FDA compounds and PeruNPDB. B. Lipinski’s five rules from the FDA and PeruNPDB data sets. C. Visual 3D representation of the chemical space as PCAs from the FDA and PeruNPDB data sets. D. Representation t-SNE of FDA and PeruNPDB data sets.

## Discussion

Peru has exceptionally high biodiversity, with numerous endemic species of mammals, reptiles, amphibians, flowering plants, and ferns, which is why has been described as a ‘megadiverse’ country [64,65], but worldwide hotspot analysis for potential conflict between food security and biodiversity conservation points out Peru as a region that is especially at risk of biodiversity loss due to agricultural expansion [66]. Thus, the conservancy of biodiversity can be considered important since historically NPs have played a key role in drug discovery, especially for illnesses such as cancer, cardiovascular and infectious diseases [67], while the growing interest in NPs and their application is evidenced by a growth of the number of published databases of NPs, and collections of structures from various organisms, geographical locations, targeted diseases, and traditional applications [68].

Currently, several NPs or NPs-derived molecules are employed in the treatment of distinct diseases, such as the antibiotic penicillin originally obtained from the fungi *Penicillium* spp [69]; the analgesic aspirin, which is the most used drug in the world, derived from salicin extracted from the bark of the willow trees *Salix alba* [70]; and the immunosuppressant tacrolimus employed in the prevention of the rejection organ after transplants, obtained from bacteria *Streptomyces tsukubaensis* [71], are some examples. Besides, NPs and their derivatives have been considered promising options to improve treatment efficiency in cancer patients and decrease adverse reactions [72], whereas vinca alkaloids [73], taxane diterpenoids [74], camptothecin derivatives [75], and epipodophyllotoxin [76], are NPs-derived anticancer compounds clinically used as chemotherapeutics; while an example of the importance of biodiversity conservation is exemplified by the tree *Taxus brevifolia*, from which the chemotherapeutic drug paclitaxel was originally extracted, that was put on the list of endangered species [77,78].

According to the data, there are fewer compounds identified in the PeruNPDB than in AfroDB, BIOFAQUIM, and NuBBE_DB_, but the chemical diversity is also higher. Of the 280 compounds characterized, 95% came from plant sources, and 5% came from animal sources. But in the BIOFACQUIM and NuBBE databases as well as plant sources, compounds derived from fungi, propolis, bacteria, and marine organisms are also described. Furthermore, the Peruvian marine biodiversity hotspot located on the northern coast has been predicted to hold 501 species, 270 genera, and 193 families [79], as marine natural products have shown an interesting array of diverse and novel chemical structures with potent biological activities [80], which includes: Cephalosporin C an antibiotic derived from marine fungi *Cephalosporium* [81], Eribulin an anticancer drug derived from halichondrin B from the natural Japanese marine sponge *Halichondria okada* [82] and the antiviral, isolated from sponge *Tethya crypta*, nucleoside Ara-A [83]. Also, Peru is considered a diverse country that has a very broad microbial diversity richness, however, remains slightly studied and exploited [84,85]. Fungi, the eukaryotic microorganisms, produce a tremendous number of NPs with diverse chemical structures and biological activities [86], such as lovastatin, the first statin approved as a hypercholesterolemic medication by the FDA, most frequently produced by *Aspergillus terreus* [87], and cyclosporine A, a potent immunosuppressant that was initially used to prevent organ rejection, isolated from the fungal species *Tolypocladium inflatum* gams [88]. Besides that no current drug has been developed from propolis, it is considered a very rich and complex chemical composition, while about 300 different chemicals components isolated from it, and which composition fluctuates according to parameters such as plant source, seasons harvesting, geography, type of bee flora, climate changes, and honeybee species [89,90]; highlighting Artepillin C, extracted from Brazilian green propolis, that showed *in vitro* [91] and *in vivo* [92]anti-inflammatory potential. These emphasize the urgency to promote and enhance the study of Peruvian NPs quantitatively and qualitatively.

Compounds from Peruvian medicinal plants have been evaluated for their antidiabetic [93], anticancer [94], antiviral [95], antibiotic [96], and antiparasitic activities [97]; however, most of the studies in the literature were *in vitro* performed over plants extracts, and little information about the potential of single compounds on these activities is described, while these promising results can be explained by synergistic interaction or multi-factorial effects between compounds present in the plant extracts studied [98]. While pharmacodynamic synergy involves multiple substances acting on various receptor targets to enhance the overall therapeutic effect, and pharmacokinetic synergy involves substances with little to no activity helping the main active principle to reach the target by improving bioavailability or by reducing metabolism and excretion, this type of assay can hide the true potential of single molecules activity between different constituents of plant extracts. [99]. Thus, the concerted effort of experimental NPs research with CADD is continuously increasing; and recently, NPs from the Peruvian native plants *Smallanthus sonchilofolius*, *Lepidium meyenii* (40 compounds) [38]; and *Uncaria tomentosa* (26 compounds) [100] were in silico analyzed for their antiviral activity against SARS-Cov-2. Also, the *in silico* polypharmaceutical potential of 84 NPs from *S. sonchifolius*, *L. meyenii*, *Croton lechleri*, *U. tomentosa*, *Minthostachys mollis*, and *Physalis peruvianus* was analyzed against Alzheimer’s disease [101].

## Conclusion

Here we present the first version of PeruNPDB, a compound database of NPs from Peru that includes 280 compounds from plant and animal sources. PeruNPDB was constructed curated, and maintained by the Computational Biology and Chemistry Research Group from the Universidad Catolica de Santa Maria, and it is freely accessible through the website https://perunpdb.com.pe/. The PeruNPDB was envisioned as a tool for virtual screening, identifying promising compounds, serving as a springboard for further biotechnological products, and providing suggestions for conservation policies. The chemoinformatic characterization and analysis of the coverage and diversity of PeruNPDB in chemical space suggest broad coverage, overlapping with regions in the drug-like chemical space. The database contains an identification code (ID), the chemical name, bibliographic reference (name of the journal, year of publication, and DOI number), kingdom, genus, and species of the natural product, SMILES notation, and classification of the natural product. As a perspective, we aim to release the second version of the PeruNPDB, enhancing and retaining its web-based user interface while adding data on biological activities and new compounds from different taxonomic ranks not included in the current version.

## Author Contributions

Conceptualization: J. L. M. F. and M. A. C-F.; data curation: H. L. B-C., L. G. R., and M. A. C-F.; formal analysis: J. L. M. F. and M. A. C-F.; funding acquisition: G. D. D-C. and M. A. C-F.; investigation H. L. B-C., L. G. R., M. A. C-P., E. G. C-R., A. E. C-L.; methodology: J. L. M. F. and M. A. C-F.; writing—review & editing: H. L. B-C., L. G. R., G. D. D-C., J. L. M. F., and M. A. C-F. All authors have read and agreed to the published version of the manuscript.

## Institutional Review Board Statement

Not applicable.

## Informed Consent Statement

Not applicable.

## Data Availability Statement

Not applicable.

## Acknowledgments

Not applicable.

